# Shedding light on Amazonian phylogeographic patterns and evolutionary history of night monkeys (Genus: *Aotus*) using reconstructed fecal metagenomic shotgun sequencing

**DOI:** 10.1101/2025.11.03.686292

**Authors:** William D. Helenbrook

## Abstract

Taxonomic classification, phylogenetics, and evolutionary history of night monkeys (*Aotus*) remain subjects of ongoing debate, particularly regarding species boundaries and the complex interplay of drivers underlying Neotropical primate diversification, including the historical role of the Amazon River as a biogeographic divide, Andean uplift, and Pliocene–Pleistocene climatic fluctuations. This study refines our understanding of *Aotus* systematics by employing fecal metagenomics to generate complete mitochondrial genomes from strategically collected samples across the lower Amazon Basin. Phylogenetic analyses integrating Maximum Likelihood, Bayesian inference, and species delimitation frameworks revealed several corrections to previously proposed biogeographic ranges and clarified evolutionary relationships among taxa. Phylogenomic reconstruction based on complete mitogenomes supports a reorganized classification into three principal clades—northern, western, and southern— originating in the Early Pliocene. Additionally, an expanded mitochondrial cytochrome *c* oxidase subunit I (COI) dataset provided greater resolution of haplogroups within multiple species, revealing fine-scale geographic structure corresponding to major river barriers. Divergence time estimates were consistent with earlier studies, indicating a most recent common ancestor of the family Aotidae at approximately 17.2 million years ago and rapid diversification within *Aotus* beginning around 4.5 million years ago. Collectively, these results refine species distributions, illuminate the evolutionary history of *Aotus*, and provide an updated phylogeographic framework with direct implications for taxonomy and conservation management across the genus.

Graphical illustration
Phylogenetic relationships among *Aotus* taxa inferred from combined complete mitochondrial genomes and single-gene COI sequences using Bayesian Inference (BI). Branches are colored according to biogeographically defined taxonomic groups. Both datasets corroborate the recovery of three major clades (i.e., northern, western, and southern) and reveal fine-scale haplogroup structure corresponding to major Amazonian river barriers. Posterior probabilities are indicated at key nodes; only values <0.99 are shown. This integrated analysis highlights concordance between full mitogenomic and single-gene datasets, clarifying species boundaries and evolutionary relationships across the genus.

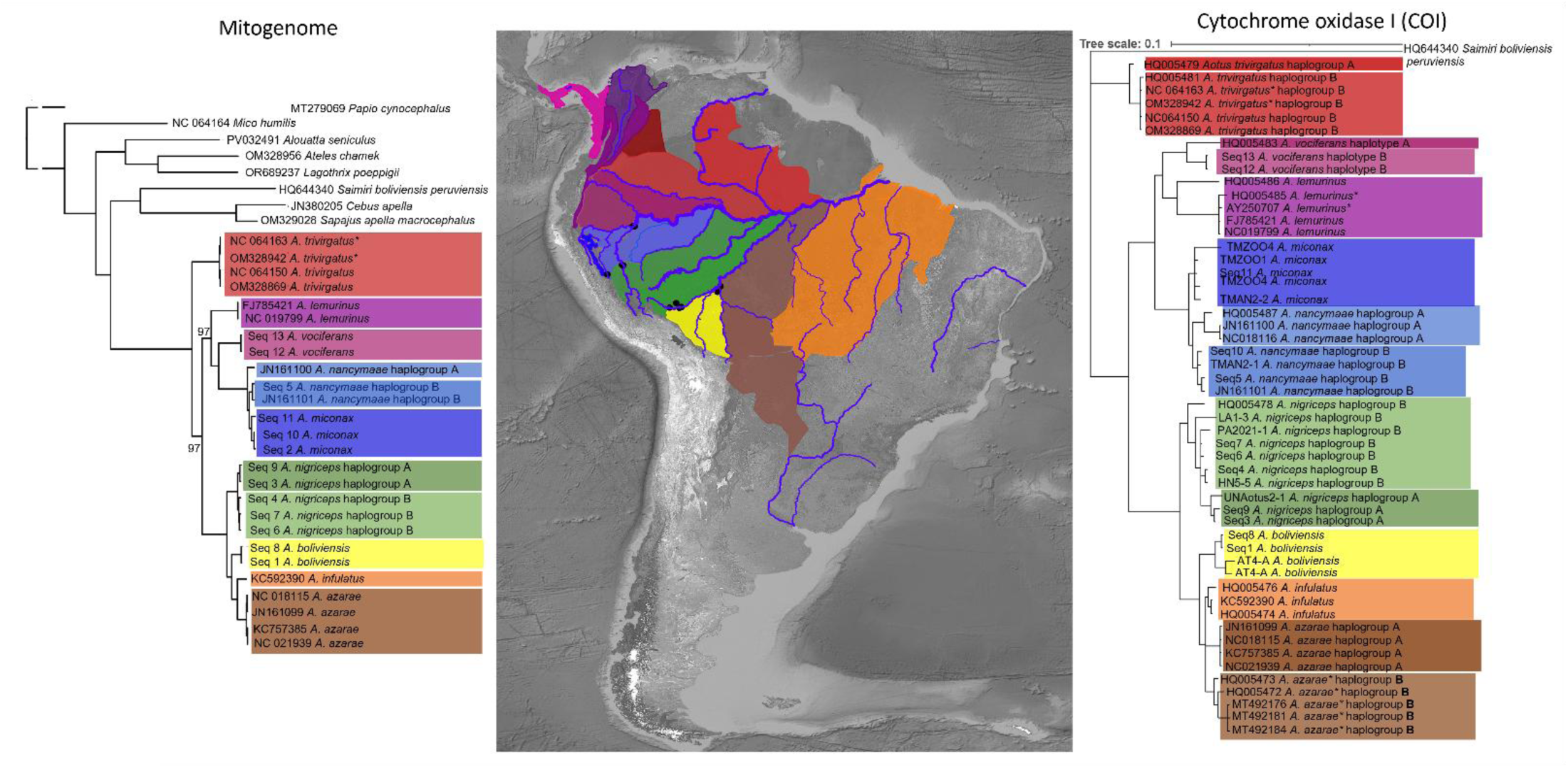

**Highlights:**

- Complete *Aotus* mitochondrial genomes were reconstructed from fecal samples across Amazon.
- Expanded COI analyses revealed haplogroups structured by rivers and areas of endemism.
- Misidentified sequences were corrected, clarifying *A. boliviensis and A. trivirgatus*.
- Biogeographic ranges were corrected, showing reduced distributions with conservation impact.

## 1. Introduction

### 1.1 Taxonomic history of Aotus

The only nocturnal monkey (Genus: *Aotus*) goes by many names, including night and owl monkeys, douroucouli to represent the French pronunciation of *Aotus* as interpreted from the Marabitanas Native Americans (Hershkovitz 1983), *macaco de noite* in Portuguese, tuta mono in Quechua-Spanish, and musmuqui (personal observation across north and south Peruvian Amazon). The taxonomic history of night monkeys has been a subject of ongoing debate and complexity for several decades (Table 1). This is partly due to relatively limited sampling efforts across much of their range, even though the genus is distributed from Panama to Argentina. Varying methodologies, mistaken origins, minimal phenotypic variance, and conflicting Amazonian geological history have contributed to challenges in classifying and understanding origin and diversity of this genus (Ashley and Vaughn, 1995; Defler and Bueno, 2007; Plautz et al., 2009; Menezes et al., 2010, Monsalve and Defler, 2010; Babb et al., 2011; Martins-Junior et al., 2022).

**Table 1:**
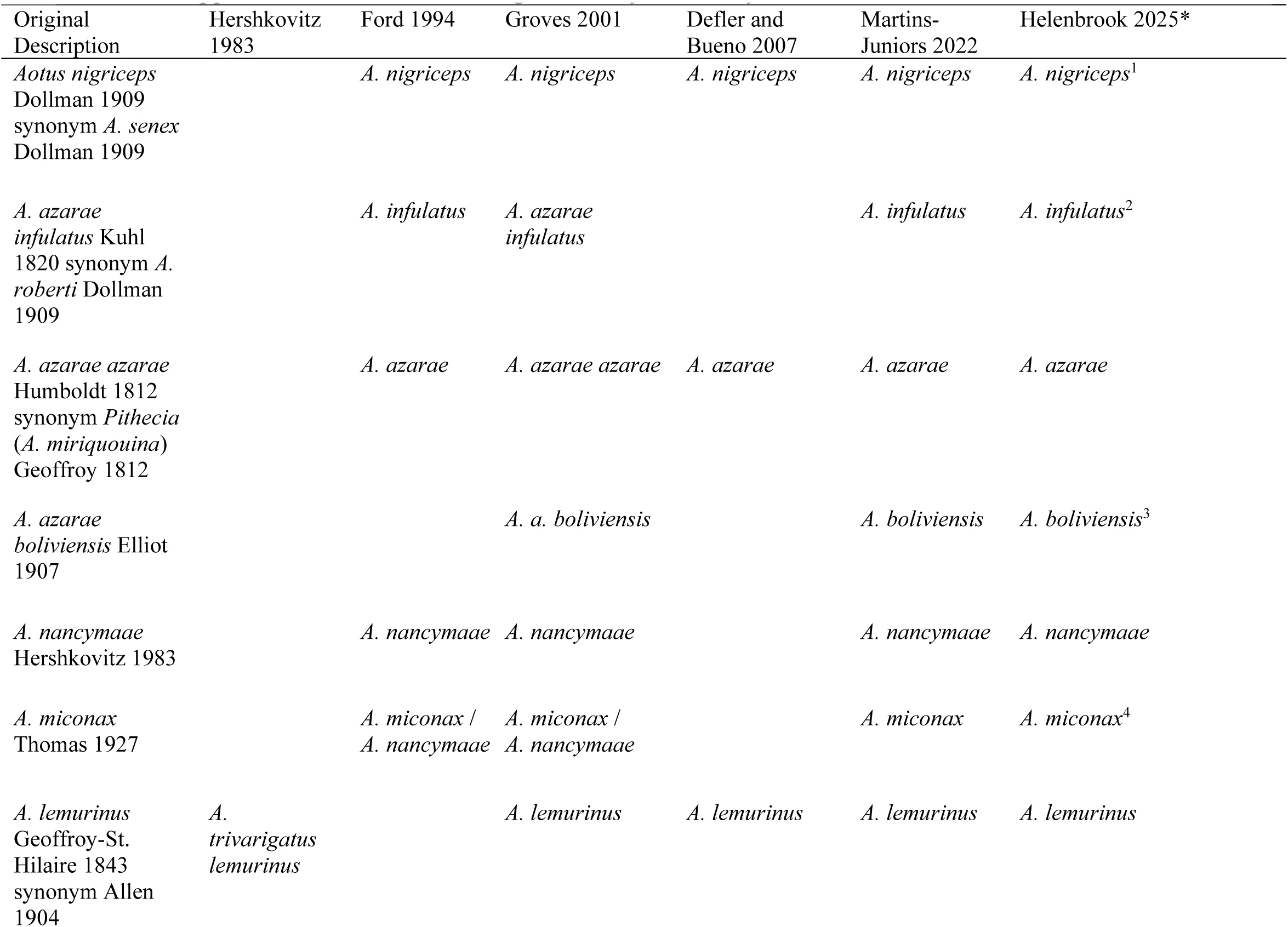
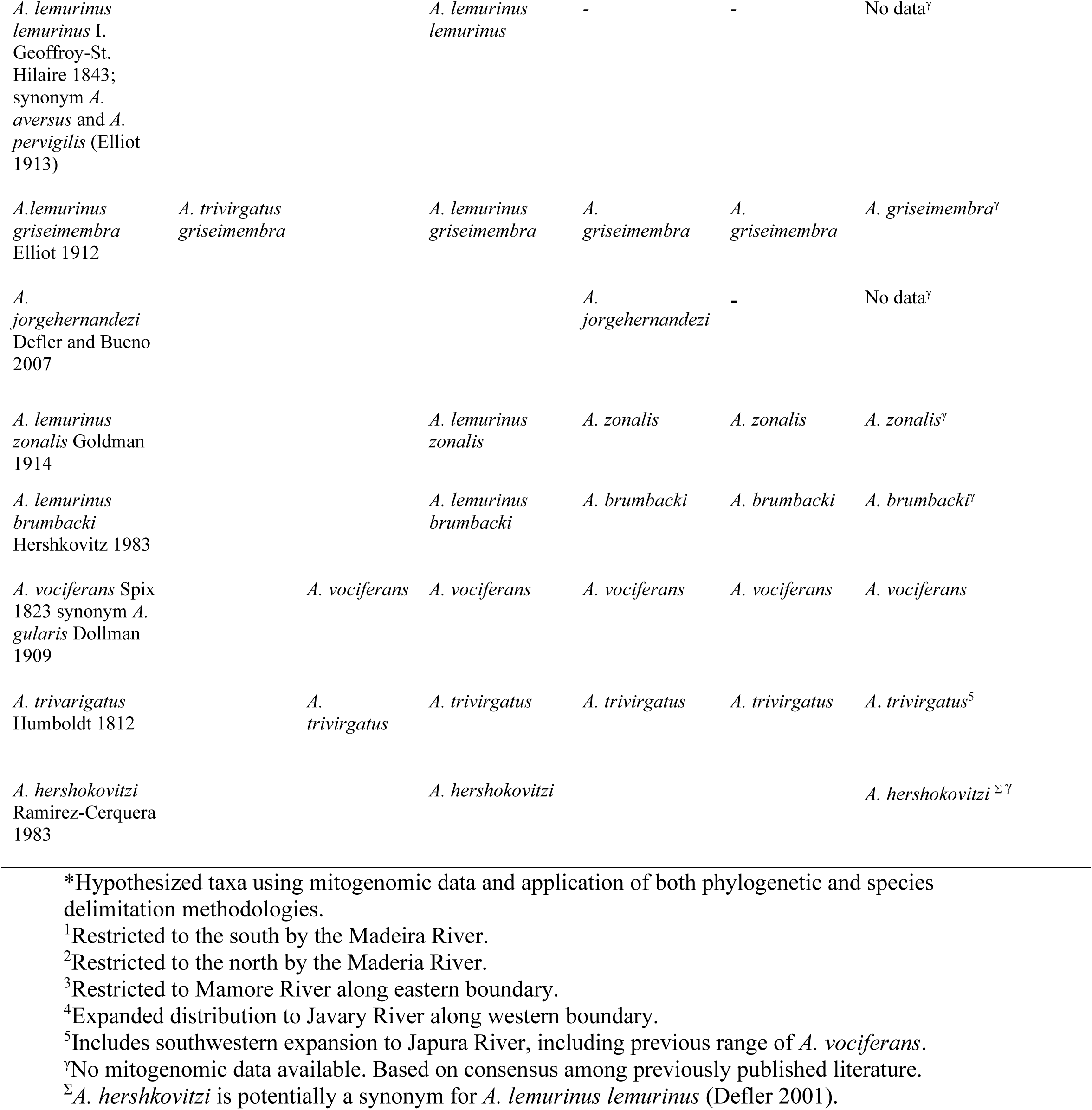
Supported classifications for night monkey taxonomy based on select studies.

The inclusion of various taxonomic methods used over the years, such as morphological descriptions of pelage color and pattern, and morphology (Hershkovitz 1983; Groves 2001), chromosomal variability (Brumback et al., 1971; Ma et al. 1985; Pieczarka et al. 1993; Torres et al. 1998; Defler and Bueno 2007) and malaria susceptibility (Hershkovitz 1983) demonstrates the extensive efforts to refine *Aotus* taxonomy. However, it also indicates the lack of consensus utilizing various methods.

As of Hershkovitz (1983), nine species were recognized based on karyotype, color, and pelage patterns. This arrangement included four southern (or red-necked) species (i., *A. azarae* including subspecies *azarae* and *boliviensis*, *A. miconax*, *A. nancymai*, *A. infulatus*) as well as four northern, gray-necked species (i.e., *A. brumbacki*, *A. lemurinus* including subspecies *lemurinus* and *griseimembra*, *A. trivirgatus*, and *A. vociferans*). There have been more recent reassessments of taxonomic diversity and status (Table 1). Molecular phylogenetic studies in *Aotus* species have focused primarily on single mitochondrial genes (Ashley and Vaughn 1995; Plautz et al. 2009; Menezes et al. 2010), and nuclear gene fragments such as Single Nucleotide Polymorphisms (SNPs) (Martins-Junior et al. 2022).

Recently, Martins-Junior et al. (2022) revisited evolutionary history descriptions of the genus and reassessed taxonomic classification. Their study suggested *A. nancymaae*, which was previously grouped with the red-necked night monkeys, is more closely related to *A. vociferans* based on select mitochondrial markers. This finding was not supported by their nuclear dataset though; however, such discrepancies are not uncommon as mitochondrial phylogenies solely represent matrilineal relationships, which may not align with the species phylogeny derived from nuclear genomic data.

Several factors indicate the possibility of further taxonomic diversity within *Aotus*, emphasizing the need for continued phylogenetic investigation. Firstly, rivers have played a significant role in impeding gene flow and contributing to allopatric speciation among *Aotus* (Helenbrook and Valdez 2025), other Neotropical primates (e.g., Ayres and Clutton-Brock 1992; Cortés-Ortiz et al. 2003; Alfaro et al. 2015; Ruiz-Garcia et al. 2017; Martins-Junior et al. 2018; Boubli et al. 2019), and an extensive list of other sympatric flora and fauna such as plants (Nazareno et al., 2019), amphibians (Moraes et al., 2016; Figueiredo-Vázquez et al., 2021), and birds (e.g., Hayes & Sewlal, 2004; Fernandes et al., 2012; Ribas et al., 2012; Pomara et al., 2014; Naka & Brumfield, 2018; Kopuchian et al., 2020). This phenomenon has resulted in allopatric speciation, particularly in regions of the Amazon basin where large rivers have acted as formidable barriers, hindering the movement of wildlife (e.g., Ayres and Clutton-Brock 1992; Ribas et al. 2012; Musher et al. 2022). Furthermore, *Aotus* taxa have been shown to exhibit the highest degree of molecular divergence in relation to river width compared to all other Platyrrhini genera (Helenbrook and Valdez 2025).

### 1.2 Metagenomics and phylogenomics

Recent advancements in metagenomics and bioinformatics have enabled the reconstruction of host genomes from diverse environmental samples, making it a valuable tool for investigating phylogenomics and evolutionary history of wildlife (Ang et al., 2020; van der Valk et al. 2020; Murchie et al., 2022). Mitogenomes can be reconstructed from fecal samples collected during routine field surveys (Srivathsan et al. 2016), providing valuable insights into species identification, genetic diversity, and evolutionary relationships, especially for data-deficient taxa that require reassessment of species boundaries and distributions. Mitogenome sequencing can be a valuable tool for resolving taxonomic uncertainties, identifying cryptic species, and providing insights into the evolutionary history and genetic diversity of species, particularly for those that are difficult to study using traditional taxonomy (Hunter et al., 2018; Davies et al., 2020).

Previous mitogenomic phylogenetic analyses of *Aotus* have been limited to a small subset of currently described taxa, and further exploration is necessary to address the taxonomic uncertainties and test hypotheses related to river-restricted dispersal. Therefore, this study aimed to expand the sampling of *Aotus* species across the Amazon basin, focusing on adjacent riverbanks of major river systems to infer host taxonomy, assess phylogenetic relationships using expanded mitogenomic datasets, and estimate divergence times based on improved phylogenomic inference and recent improvements to our understanding of geological history in the region. The study hypothesizes that *Aotus* taxonomy in the Amazon basin requires additional revision, where further taxonomic refinement exists, driven in large part by restricted gene flow across several major river barriers.

## 2. Methods

Fecal samples were non-invasively collected from fifteen (N = 15) night monkey individuals representing both currently recognized and hypothesized taxa across strategic sites in Brazil, Bolivia, and Peru, located near major river barriers and spanning distinct ecoregions. These samples were used to generate complete mitochondrial genomes; however, only thirteen yielded sequences with sufficient quality and coverage for inclusion in subsequent phylogenomic analyses (Figure 1). *Aotus* groups were surveyed on established trails in the evening and early morning hours. Fecal samples were gathered using plastic netting strategically positioned beneath nests or frequently traversed areas to ensure avoidance of soil contamination and disturbance from other animals. Nets were examined within twelve hours at least twice a day, and geospatial data recorded. Seven additional *Aotus* mitochondrial genomes with known provenance were retrieved from the National Center for Biotechnology Information (NCBI) database.

**Figure 1:**
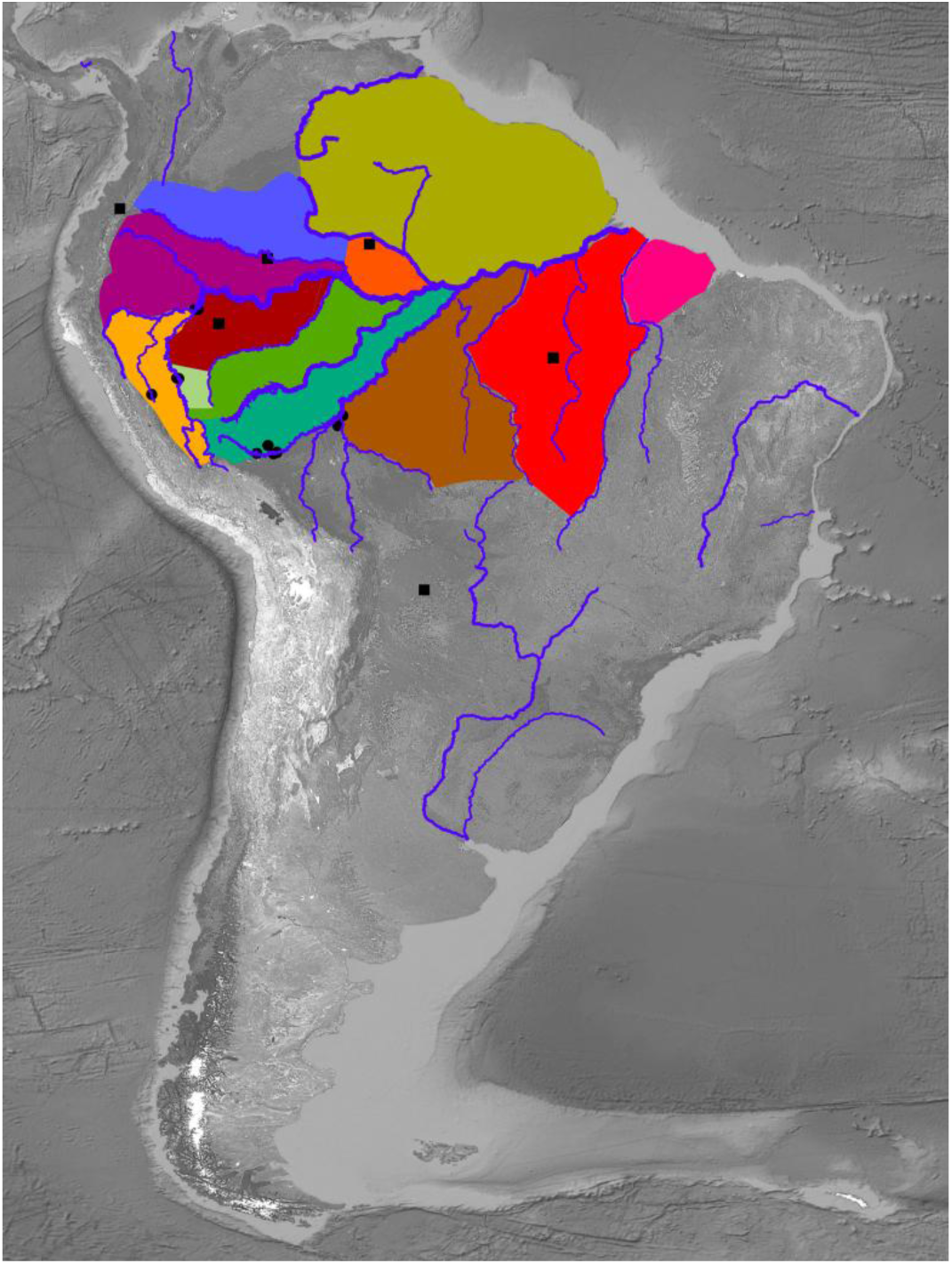
Areas of endemism were delineated based on hypotheses generated from Helenbrook and Valdez (2025). Field sites (indicated by circles) were selected within the Amazon Basin at boundaries largely defined by river barriers. Squares represent mitochondrial genomes from previous studies with verified provenance.

An additional dataset of cytochrome oxidase I (COI) sequences (N = 11; Figure 5) was independently amplified from a separate collection of fecal samples using gene-targeted PCR primers previously described by Helenbrook and Whipps (2021). These sequences were integrated with homologous COI regions extracted from the mitogenomes assembled in this study (N=13) and also previously published COI genes (N = 29) retrieved from NCBI. These single gene sequences were analyzed using the same phylogenetic framework as the full mitogenome dataset to evaluate whether the broader set of *Aotus* haplotypes corroborated mitogenomic relationships or provided additional special resolution. Published mitogenomes without geographically demarcated locations were not included. All fieldwork was conducted in accordance with the legal requirements of Peru and Brazil, and with the consent of private landowners.

### 2.1 DNA extraction and initial screening

Samples were stored in RNAlater (Qiagen Inc., Valencia, CA) and DNA was extracted using Zymo Quick-DNA Fecal/Soil Kit (Zymo Research, Irvine, CA) following the manufacturer’s instructions. Partial cytochrome c oxidase subunit II (COII) sequence were amplified in both directions and evaluated by agarose gel electrophoresis to ensure host species prior to genomic analysis using previously published primers for *Aotus* (Helenbrook and Whipps 2021). DNA was amplified by PCR in 50 μL reaction volumes in Taq 2× Master Mix New England Biolabs, Ipswich, MA, 0.25 μM of each primer, and 3 μL of template DNA using an C1000 Thermal Cycler BioRad for 40 cycles at 95 °C for 30s, 53 °C for 60 s, and 68 °C for 60 s, preceded by an initial denaturation at 95 °C for 3 min, and followed by a final extension at 68 °C for 7 min. DNA Analysis Facility at Yale ran sequences using the ABI BigDye Terminator Cycle Sequencing Ready Reaction Kit v3.1, on the 3730xl DNA Analyzer Applied Biosystems, Thermo Fischer Scientific, Inc. with a 96-capillary 50cm array. Sequences were screened in FinchTV 1.4.0 and edited in JalView to identify base-calling errors.

Shotgun metagenomics was then performed using Illumina high throughput DNA sequencing (NovaSeq 2x150bp paired-end). Libraries were pooled based on effective concentration and required sequencing data volume. Starting material (100 ∼ 1000ng) passed quality control. DNA was fragmented by acoustic disruption using Covaris S220 and then end repair, dA-tailing, adapter ligation and purification. The preliminary quantification and dilution of the library was performed using Qubit3.0, and then Agilent 2100 was used to determine the insert size and nucleic acid concentration of the resulting library sample. Effective concentration of each sample library in mixture was determined by qPCR before sequencing to ensure accuracy of sample concentration and reliability of sequencing data. Quality assessment covered data volume, sequence quality distribution, base distribution, sequencing error rate, data filtering, exogenous contamination, etc. Base calling was achieved with built-in Real-Time Analysis software which performed conversion of four fluorescent signals obtained from charge-coupled device to binary base call (BCL) data. BCL data were then converted to FASTQ files using bcl2fastq (v2.17, part of the software package provided by Illumina). Data demultiplexing was performed simultaneously based on index information. Primary analysis was performed using built-in software HiSeq Control Software. Reads of no more than 2 out of the 25 cycles with chastity values below 0.6, were called PF (Pass Filter). PF clusters were converted by bcl2fastq and stored in FASTQ format. Error rate distribution was used to spot high error rates in middle and towards end reads. Base quality scores for Illumina platforms were expressed in Q Phred, calculated from error rate (e) using the formula Qphred = -10log10(e). Raw data were filtered based on the following criteria: (1) remove pair-end (PE) reads containing adapter sequences; (2) remove PE reads with Q scores below 20 for over 50% of the entire sequence; (3) remove PE reads with N composition greater than 10%.

### 2.2 Bioinformatics

Mitogenomes were reconstructed from multiple short reads generated from sequencing in which a reference mitogenome and iterative mapping and assembly are used to generate a complete mitogenome assembly. Raw paired-end files were first trimmed to remove adapter sequences and low-quality sequences using Trim Galore! (Krueger 2015). New mitochondrial reference genomes were mapped for each *Aotus* species using Bowtie2 (Version 2.5.4; Langmead and Salzberg 2012). Each sample was mapped against available *Aotus nancymaae* reference genomes (NC_018116). Assembly and error correction performed using SPADES 3.15.3 (Python 3.9.0; Bankevich et al. 2012). Coverage of the mitochondrial genome was estimated using the "plotCoverage" function (Ramírez et al. 2016). All mitogenomes were annotated using MITOS2, analyzing all protein-coding genes, as well as tRNA and rRNA genes (Donath et al. 2019). To avoid nuclear DNA sequences of mitochondrial origin (numts), all coding regions were annotated for open reading frames and stop codons. All generated sequences were manually analyzed prior to GenBank submission.

### 2.3 Sequence alignment

Newly reconstructed mitogenomes were aligned with select Platyrrhini mitogenomes from GenBank (Supplemental Table 1) and annotated for BankIt Submission at the National Library of Medicine. Only unique *Aotus* mitogenomes with verified origin in published literature were included. A total of twenty-six *Aotus* mitogenomes were analyzed, including thirteen complete mitogenomes generated in this study. Sequences were aligned using MAFFT (version 7.407_1) set with 0.123 gap extended penalty and 1.53 gap opening penalty and curated with BMGE (Block Mapping and Gathering with Entropy) using PAM matrix 250, maximum entropy threshold of 0.5, gap rate cut-off of 0.5, and minimum block size of 5 (Criscuolo et al. 2010). A final alignment of 15423 bp was used due to low coverage of the last tRNA gene in several samples. In total, there were eight Platyrrhini primate genera including representatives from five currently recognized *Aotus* species, and one Cercopithecoidea (*Papio papio*) used as an outgroup.

### 2.4 Phylogenetic inference

Thirty-three Platyrrhini mitogenomes (N=33) of known provenance were used to reconstruct phylogenetic trees using Maximum-Likelihood (ML) and Bayesian Inference (BI) methods. Best-fit substitution model was selected using both MEGA 11 (Tamura et al. 2021) and Smart Model Selection 1.8.1_1 in PhyML (version 1.8.1_1; Lefort et al. 2017). A generalized time reversible model with gamma-distributed rate variation and a proportion of invariant sites (GTR + G + I) was selected as the best model under Akaike information criterion, Bayesian information criterion, and decision theoretic frameworks. I applied the Shimodara-Hasegawa-like approximate likelihood ratio test (SH-aLRT) with 10,000 replicates and non-parametric bootstrap with 1000 replicates to assess branch support (Anisimova et al. 2011). Bayesian phylogenetic inference analysis was inferred with Mr. Bayes (version 3.2.7_0; Huelsenbeck et al. 2005), utilizing the same generalized time reversible substitution model (GTR + G + I) with three data partitions within alignment (codon positions 1, 2, and 3). Four independent Markov Chain Monte Carlo (MCMC) runs were performed, with 4 chains, 1.25 million generations, sampling every 1000 generations. An initial twenty percent of sampled data were discarded as burn-in for each run. The single-gene COI dataset was analyzed using the same methodological framework as the complete mitogenome analysis.

### 2.5 Species delimitation

Computational species delimitation (CSD) algorithms, capable of delineating or confirming evolutionary significant units (ESUs), are now integrated as an additional taxonomic approach (Moritz 1994). These algorithms operate based on the phylogenetic species concept outlined by Eldredge and Cracraft (1980). Curated sequences were used in tree-based species delimitation analyses (i.e., Bayesian Poisson Tree Processes, bPTP) (Zhang et al. 2013). The resulting trees were subjected to PTP-based species delimitation after excluding outgroups. PTP is a probabilistic model-based method used to infer species boundaries directly from molecular phylogenies without relying on predefined genetic distance thresholds or morphological criteria. PTP assigns each branch in the tree to either an intra- or interspecific category.

### 2.6 Time-calibration of phylogenetic tree

Estimated divergence dates and associated confidence intervals of Platyrrhini species were inferred in MEGA11 with Timetree, inferred by applying the RelTime method with user-supplied phylogenetic tree with branch lengths (Tamura et al. 2018; Mello 2018). I constrained tree topology based on branching patterns that had > 90% bootstrap support in the maximum likelihood phylogenetic analyses and posterior clade credibilities of 1.0 with Bayesian analyses. Minimum and maximum time boundaries on nodes were based on minimum and maximum calibration TimeTree 5 calibration constraints, which including fossil evidence, geological events, biogeographical data, paleoclimatic and molecular studies (Ho et al. 2015; Kumar et al. 2022). Times were not estimated for outgroup nodes because the Reltime method uses evolutionary rates from the ingroup to calculate divergence times and does not assume that evolutionary rates in the ingroup clade apply to the outgroup. A discrete Gamma distribution was used to model evolutionary rate differences among sites (5 categories (+G, parameter = 0.1935)). The rate variation model allowed for some sites to be evolutionarily invariable ([+I], 49.85% sites). Codon positions 1, 2, and 3 were included. The final dataset comprised 16,571 nucleotide positions, with all sites containing gaps or missing data excluded from the analysis (complete deletion option).

## 3. Results

### 3.1 Metagenomic analysis

Illumina sequencing of fecal metagenomes yielded an average of 42,618,489 million reads per sample (range: 34.3 – 52.5 million) with an average sequencing depth of 26.8x (Table 2). Thirteen *Aotus* mitogenomes were assembled from eleven disparate field sites in Peru, Bolivia and Brazil (Figures 1). All protein-coding genes were correctly translated for full mitogenomes without any premature stop codons (available through NCBI), indicating that no nuclear mitochondrial-like sequences (numts) are present. Reconstructed mitogenomes included twenty-one (N=21) transfer RNA genes, two (N=2) ribosomal RNA genes, thirteen protein coding genes and the control region. All assembled mitogenomic sequences were novel and an additional thirteen previously published mitogenomes with known provenance were included. Two additional samples, not included here, showed low average mitochondrial genome coverage and were excluded from further analysis. The first complete mitochondrial genomes are reported for *Aotus nigriceps*, *A. vociferans*, and *A. boliviensis*, with evidence indicating that two previously published sequences were likely misidentified (i.e., NC_064163 and OM328942).

**Table 2:** Mitogenomic samples, metagenomic results, and associated taxonomic information generated using different data sets and analysis. It should be noted that the *A. boliviensis* samples reported here represent a previously undocumented geographic area and should not be confused with specimens previously assigned to the State of Rondônia, Brazil.

| Sample Name | Location | Sequence Length (bp) | Reads | Average Coverage Depth | Previous Taxonomic Description (IUCN) | Species Delimitation Results | Phylogenomic Species Concept<sup>b</sup> | Cytochrome Oxidase I (COI) |
| --- | --- | --- | --- | --- | --- | --- | --- | --- |
| Seq3 (UN2-1) | -8.382516°<br>-74.476300° | 15495 | 37,322,707 | 16.2x | *Aotus nigriceps* | *A. nigriceps* | *A. nigriceps*<br>haplogroup A | *A. nigriceps*<br>haplogroup A |
| Seq9 (SJ1-10) | -8.341713°<br>-74.593348° | 15529 | 47,396,409 | 29.6x | *A. nigriceps* | *A. nigriceps* | *A. nigriceps*<br>haplogroup A | *A. nigriceps*<br>haplogroup A |
| Seq6 (LA1-1) | -12.571568°<br>-70.100263° | 15634 | 41,462,943 | 11.7x | *A. nigriceps* | *A. nigriceps* | *A. nigriceps*<br>haplogroup B | *A. nigriceps*<br>haplogroup B |
| Seq4 (HN5-5) | -12.127348°<br>-69.479799° | 15569 | 46,093,947 | 28.7x | *A. nigriceps* | *A. nigriceps* | *A. nigriceps*<br>haplogroup B | *A. nigriceps*<br>haplogroup B |
| Seq7 (PA2021-1) | -12.505498°<br>-68.975557° | 15640 | 44,814,072 | 59.8x | *A. nigriceps* | *A. nigriceps* | *A. nigriceps*<br>haplogroup B | *A. nigriceps*<br>haplogroup B |
| Seq1 (AT4-B) | -12.522659°<br>-68.980634° | 16582 | 52,536,543 | 28.0x | *A. azarae*<br>*boliviensis* | *A. boliviensis* | *A. boliviensis* | *A. boliviensis* |
| Seq8 (PM1-2) | -12.596865°<br>-69.171650° | 14953 | 45,963,484 | 13.5x | *A. azarae*<br>*boliviensis* | *A. boliviensis* | *A. boliviensis* | *A. boliviensis* |
| Seq2 (TMAN2-1) | -9.264505°<br>-75.997702° | 15488 | 37,733,006 | 23.0x | *A. miconax* | *A. miconax* | *A. miconax* | *A. nancymaae*<br>haplogroup B |
| Seq10 (TMAN2-3) | -9.264505°<br>-75.997702° | 16469 | 35,103,545 | 32.1x | *A. miconax* | *A. miconax* | *A. miconax* | *A. nancymaae*<br>haplogroup B |
| Seq11<br>(TMZOO1) | Captive<br>population | 16469 | 41,193,436 | 34.1x | *A. miconax* | *A. miconax* | *A. miconax* | *A. miconax* |
| Seq5 (ANMA1-1) | -4.504621°<br>-73.393199° | 16450 | 48,468,167 | 33.1x | *A. nancymaae* | *A. miconax* | *A. nancymaae*<br>haplogroup B | *A. nancymaae*<br>haplogroup B |
| Seq12 (AVN1-1) | -4.508318°<br>-73.587731° | 15425 | 40,946,298 | 50.3x | *A. vociferans* | *A. vociferans* | *A. vociferans* | *A. vociferans*<br>haplogroup B |
| Seq13 (AVN1-6) | -4.508318°<br>-73.587731° | 15494 | 41,641,380 | 29.0x | *A. vociferans* | *A. vociferans* | *A. vociferans* | *A. vociferans*<br>haplogroup B |

<sup>β</sup>Using Molinari (2023) definition

### 3.2 Phylogenetic inference

Phylogenetic topologies inferred from both Maximum Likelihood (ML: Figures 2 and 4) and Bayesian Inference (BI: Figure 3) analyses were largely congruent, with the exception of *Aotus trivirgatus*. Under ML, *A. trivirgatus* occupies an intermediate phylogenetic position, whereas BI suggests it is basal to the remaining species, representing the earliest-diverging lineage. The divergence of *A. trivirgatus* from the southern clade was the only node exhibiting weak statistical support under ML. Despite this topological discrepancy, both analyses consistently recover three distinct clades within the genus, herein referred to as the northern, southern, and western groups, although additional mitogenomic data are needed for several species from the northern range (*A. zonalis*, *A. griseimembra*, and *A. brumbacki*). Pairwise genetic distances further indicate that *A. trivirgatus* is nearly equidistant from the western and southern groups, consistent with its intermediate position in the ML analysis.

**Figure 2:**
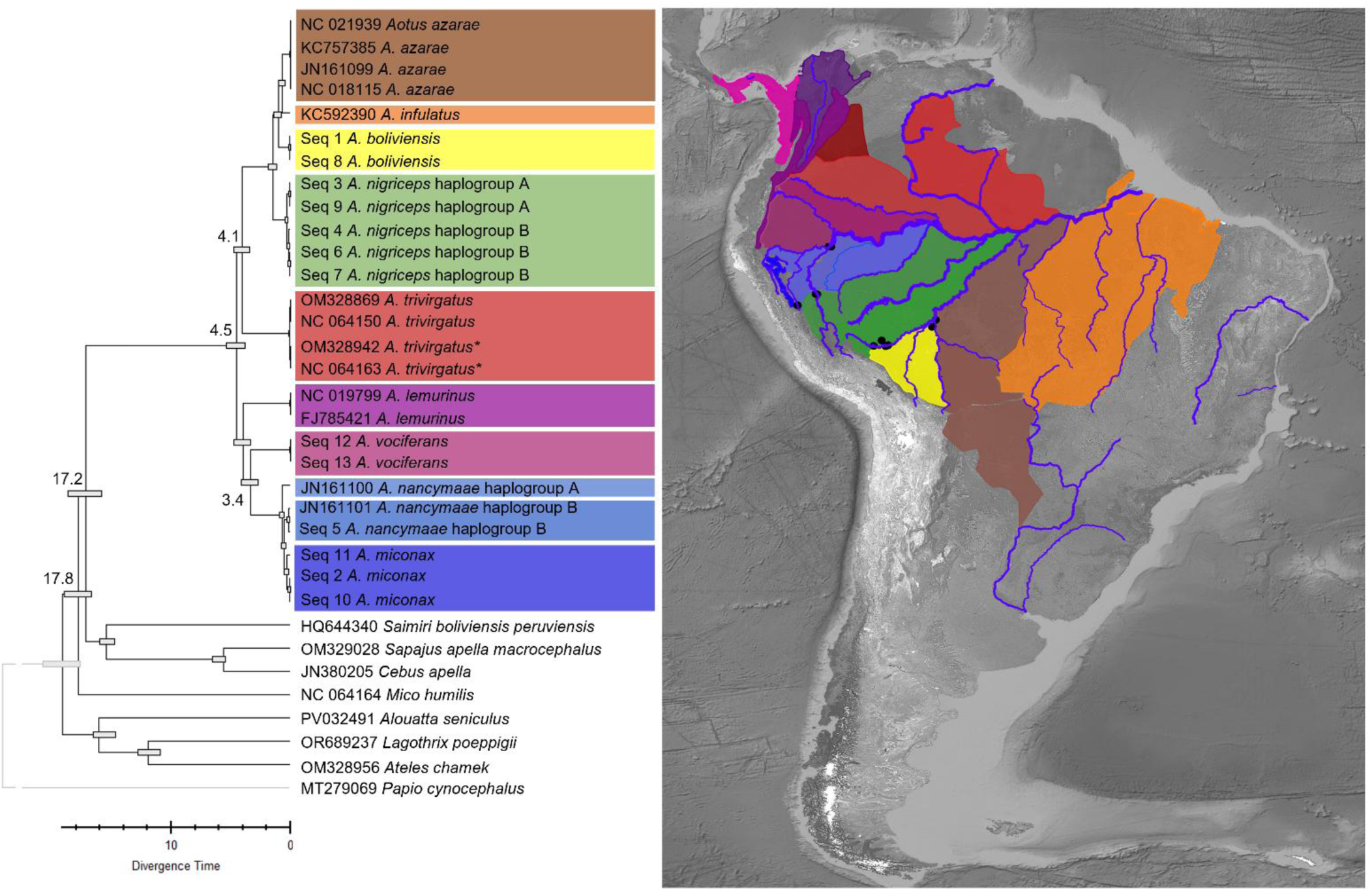
Timetree inferred using the RelTime method. The analysis used a user-provided phylogeny and the Tamura-Nei model of nucleotide substitution (Tamura and Nei, 1993), with branch lengths estimated under the Maximum Likelihood framework (Felsenstein, 1981). Site-specific evolutionary rate variation was modeled with a discrete Gamma distribution across five categories (parameter = 1.5927), and 51.85% of sites were considered invariant. Divergence times were estimated using four calibration constraints, with 95% confidence intervals indicated by bars at each node (Tamura et al., 2012; Mello et al., 2017). Times are not shown for outgroup nodes because RelTime does not assume equal evolutionary rates between ingroup and outgroup lineages (Tamura et al., 2012). The dataset comprised 34 coding sequences including 1st, 2nd, 3rd, and non-coding positions, totaling 16,570 nucleotide sites. Analyses were performed in MEGA12 using up to four parallel threads, and the estimated log likelihood of the tree was -86,365.39. * Two specimens originally labeled as *Aotus vociferans* consistently grouped with *A. trivirgatus* in both Maximum Likelihood and Bayesian analyses, indicating probable misidentification in prior datasets.

**Figure 3:**
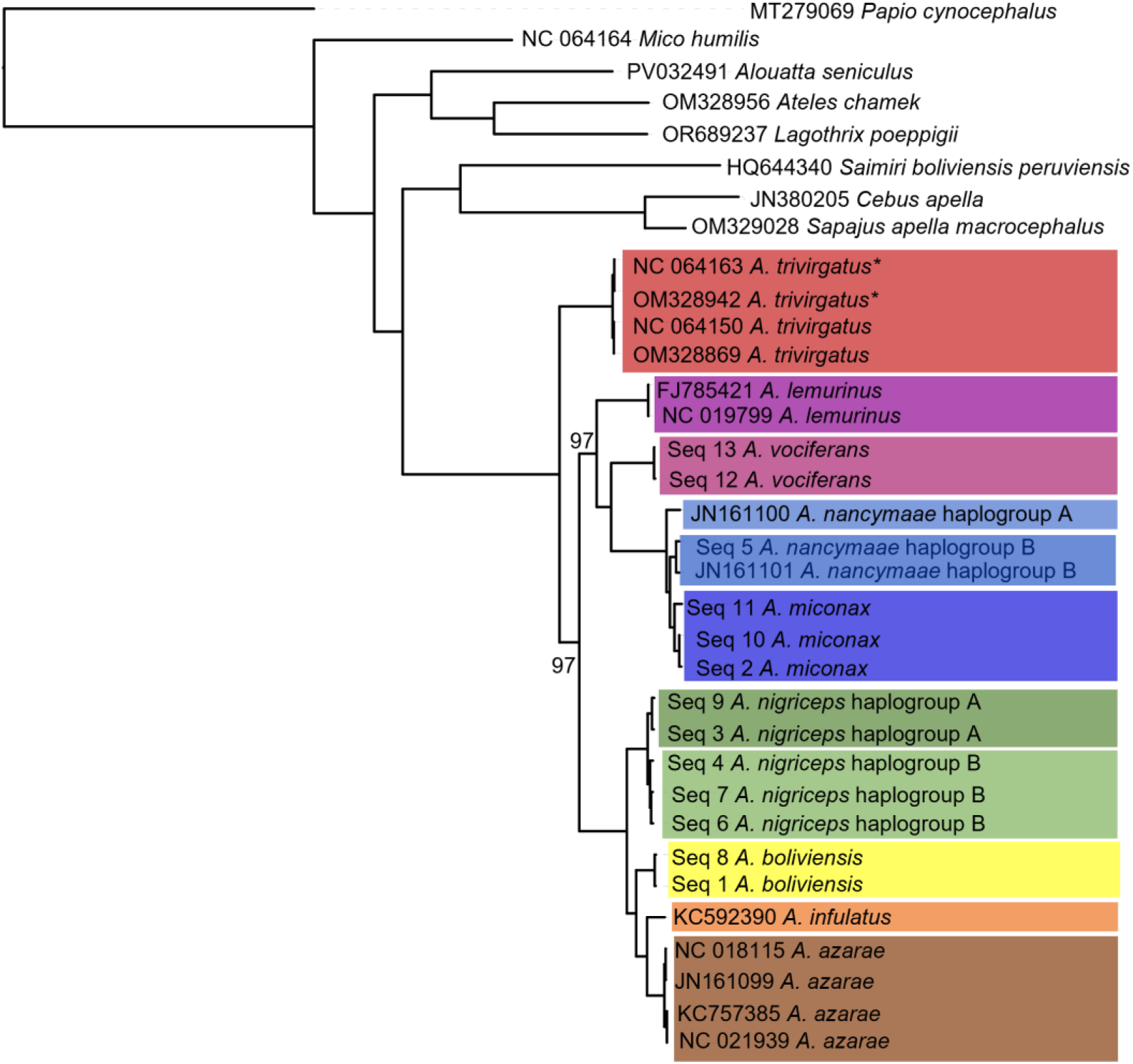
Bayesian phylogenetic tree of mitogenomes inferred using MrBayes v3.2.7. Analyses were run for 100,000 generations with four independent Markov Chain Monte Carlo (MCMC) chains per run (nrun = 4; nchain = 4), sampling every 500 generations and discarding the first 25% of samples as burn-in. The GTR substitution model (nst = 6) with equal rates across sites was applied. Convergence diagnostics and consensus calculations were performed using the sumt and sump functions. Posterior probabilities are 1 unless otherwise indicated. Branch lengths are proportional to the number of substitutions per site. *Two specimens originally labeled as *Aotus vociferans* consistently grouped with *A. trivirgatus* in both Maximum Likelihood and Bayesian analyses, indicating probable misidentification in prior datasets.

**Figure 4:**
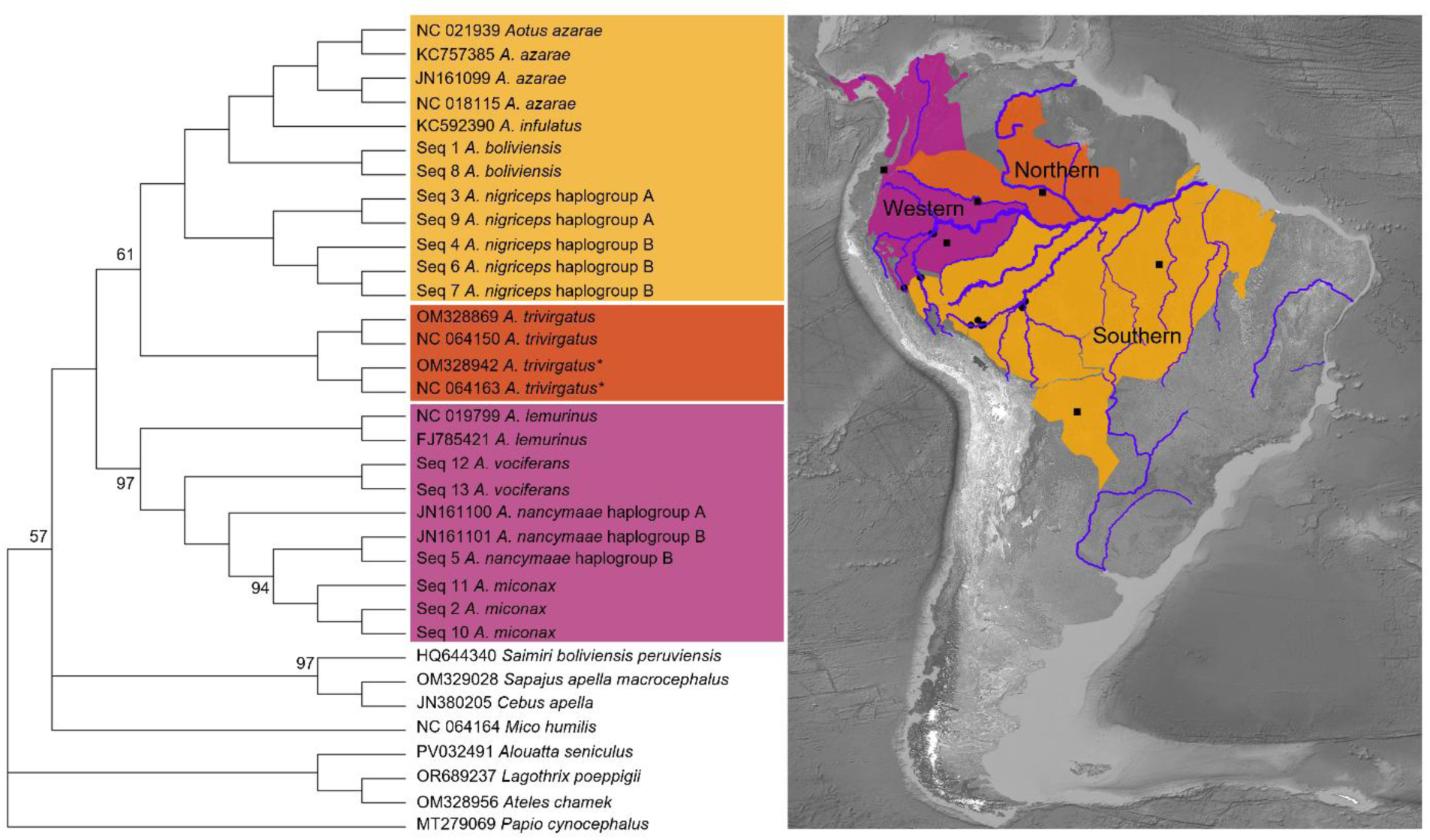
Evolutionary analysis was conducted using the Maximum Likelihood method. The bootstrap consensus tree inferred from 500 replicates (Felsenstein, 1985) represents the evolutionary history of the taxa analyzed, where branches corresponding to partitions reproduced in less than 50% of replicate trees are collapsed. Support values are greater than 0.99 unless otherwise noted. The initial tree for the heuristic search was selected by choosing the tree with the superior log-likelihood between a Neighbor-Joining (NJ) tree (Saitou and Nei, 1987) and a Maximum Parsimony (MP) tree (Kluge and Farris, 1969). The NJ tree was generated using a matrix of pairwise distances computed using the Tamura-Nei model (Tamura and Nei, 1993). The MP tree had the shortest length among 10 MP tree searches, each performed with a randomly generated starting tree. The evolutionary rate differences among sites were modeled using a discrete Gamma distribution across five categories (+G, parameter = 1.5897), with 51.83% of sites deemed evolutionarily invariant (+I). The analytical procedure encompassed 34 coding nucleotide sequences using 1st, 2nd, 3rd, and non-coding positions, with 16,570 positions in the final dataset. Evolutionary analyses were conducted in MEGA12 (Tamura et al., 2021) utilizing up to four parallel computing threads. *Two specimens originally labeled as *Aotus vociferans* consistently grouped with *A. trivirgatus* in both Maximum Likelihood and Bayesian analyses, indicating probable misidentification in prior datasets.

Across all described and available *Aotus* taxa, maximum pairwise mitochondrial divergence reached 5.4%, observed between *A. nancymaae* and *A. azarae infulatus.* Pairwise genetic distances were generally low among several currently recognized species and subspecies, suggesting limited molecular differentiation across parts of the genus (Figure 2). Notably, samples of *Aotus nancymaae* and *A. miconax*, together with an intermediate haplogroup, exhibited modest but consistent molecular divergence corresponding to their biogeographic separation. These patterns suggest the presence of three closely related genetic groups, encompassing the currently described *A. miconax* and *A. nancymaae*. However, the maximum pairwise divergence between these two described species was approximately 1%, indicating relatively shallow genetic differentiation despite their geographic isolation. A distinct mitochondrial lineage, herein referred to as the *Aotus nancymaae* haplogroup B, was recovered between the principal clades of *A. nancymaae* and *A. miconax*. This mitogenomic haplogroup exhibits intermediate divergence (∼0.8%) and is phylogenetically positioned between both species.

Both ML and BI analyses also identified two distinct haplogroups within *Aotus nigriceps*, though pairwise divergence between them was minimal, as low as 0.3% in some comparisons, suggesting very recent lineage divergence or intraspecific population structure within the species. The historically described subspecies *Aotus azarae boliviensis* and *Aotus azarae azarae* were each recovered as distinct mitochondrial haplogroups in both Maximum Likelihood (ML) and Bayesian Inference (BI) analyses. Despite clear separation, pairwise genetic distances among these taxa were low, consistent with recent divergence within the southern lineage. The smallest genetic distance was observed between *A. boliviensis* and *A. infulatus* (1.3%), while *A. azarae* exhibited a similar divergence from *A. infulatus* (1.2%).

In the northern Amazon Basin, the region between the Japurá River to the south and the Negro River to the northwest has traditionally been attributed to *Aotus vociferans*. However, two previously published mitochondrial genomes from this area (GenBank accessions NC_064163 and OM328942) exhibit the greatest molecular similarity to *A. trivirgatus*, rather than to *A. vociferans*. In contrast, newly generated mitogenomes from the southern portion of the *A. vociferans* range obtained in this study are clearly distinct, showing an average sequence divergence of 4.3% from all *A. trivirgatus* sequences. These results substantially revise the known distributions of both taxa, expanding the range of *A. trivirgatus* by approximately 205,990 km², nearly doubling its previously recognized extent, while reducing the estimated range of *A. vociferans* from 393,746 km² to 189,769 km², a decrease of 51.8%.

Species delimitation analyses using the Poisson Tree Processes (PTP) model supported the recognition of nine distinct *Aotus* species based on available mitochondrial genomes with most partitions receiving support >0.95: *A. trivirgatus*, *A. vociferans*, *A. nigriceps*, *A. boliviensis*, *A. azarae*, *A. infulatus*, and *A. nancymaae*. Two additional taxa exhibited relatively low maximum likelihood partition support (*A. miconax*, 0.40; *A. lemurinus*, 0.50). In the case of *A. miconax*, available mitogenomic sequence was derived from a single, captive individual, which clustered with samples collected from Tingo María to the eastern bank of the Ucayali River, an area previously described as part of the *A. nancymaae* range.

Phylogenetic relationships inferred from the cytochrome c oxidase subunit I (COI) gene were reconstructed using an expanded dataset consisting of sequences generated in this study from a separate set of fecal samples, combined with COI genes extracted from complete mitogenomes generated in this study, and additional sequences retrieved from GenBank with known provenance (Figure 5). The resulting tree recovered the same major topological patterns as the mitogenomic phylogeny. With the inclusion of a larger number of samples, several well-supported intraspecific haplotypes were resolved within multiple taxa. Distinct COI haplotypes were detected in *Aotus trivirgatus*, *A. vociferans*, *A. nancymaae*, *A. nigriceps*, and *A. azarae*, each corresponding to populations separated by major Amazonian river barriers.

**Figure 5:**
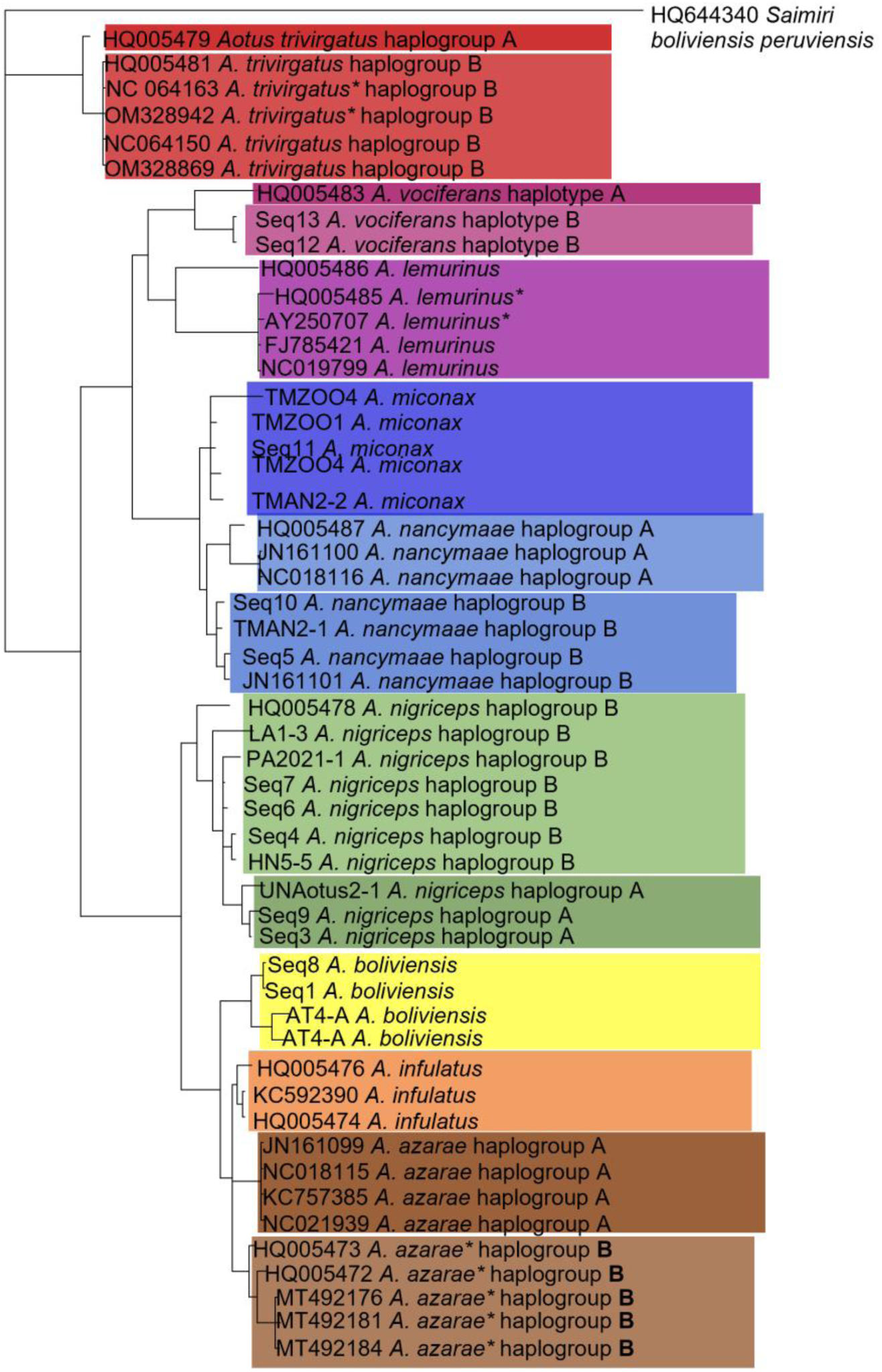
Bayesian phylogenetic tree of cytochrome oxidase I (COI gene) inferred using MrBayes v3.2.7. Analyses were run for 100,000 generations with four independent Markov Chain Monte Carlo (MCMC) chains per run (nrun = 4; nchain = 4), sampling every 500 generations and discarding the first 25% of samples as burn-in. The GTR substitution model (nst = 6) with equal rates across sites was applied. Convergence diagnostics and consensus calculations were performed using the sumt and sump functions. Posterior probabilities are 1 unless otherwise indicated. Branch lengths are proportional to the number of substitutions per site. *Indicates probable misidentification in prior datasets, and the sequences have been renamed based on BLASTN results and consistent clustering in both Maximum Likelihood and Bayesian Inference analyses.

### 3.3 Evolutionary history

Molecular dating analyses indicate that *Aotus* diverged from Cebidae, including *Saimiri*, *Sapajus*, and *Cebus*, approximately 17.2 million years ago. The monophyly of the genus was recovered with maximal support across all analyses, with a crown age estimated at 4.5 million years ago (95% Highest Posterior Density [HPD] 4.0-5.0 Ma), corresponding to the Early Pliocene. Divergence estimates among the three primary clades (i.e., northern, western, and southern) overlap temporally, reflecting a period of rapid diversification within the genus. Consistent with the topology observed in Bayesian Inference (BI), *A. trivirgatus* occupies a basal position relative to the remaining clades, whereas Maximum Likelihood (ML) analyses suggest an initial split into two lineages, which then subsequently diverge into the northern and southern groups. The divergence between *A. vociferans* and *A. nancymaae* is estimated at 3.4 Ma (95% HPD: 3.0–3.8 Ma), and *A. nigriceps* diverged from the *A. azarae* complex around 1.5 Ma (95% HPD: 1.2–1.8 Ma).

## 4. Discussion

Understanding species boundaries and evolutionary relationships within *Aotus* remains a persistent challenge in primate taxonomy. The genus is characterized by cryptic morphology, overlapping distributions, and limited comparative data across its range. Although previous studies have applied genetic and morphological tools to address these issues, debates over classification and lineage divergence continue. Many of these uncertainties stem from sparse geographic sampling and a reliance on single gene markers. This study adds to the growing literature by reconstructing complete mitochondrial genomes from fecal samples collected in previously underrepresented areas, particularly across river systems known to act as natural barriers to dispersal and gene flow. The findings provide new phylogenomic insights that clarify and challenge current taxonomic frameworks.

### 4.1 Phylogenetic insights and systematic implications

Both Maximum Likelihood (ML) and Bayesian Inference (BI) analyses recovered the monophyly of *Aotus* with maximal statistical support, consistent with previous molecular studies (Menezes et al., 2010; Martins-Junior et al., 2022). The estimated crown age of the genus, approximately 4.5 million years ago (95% HPD: 4.0–5.0 Ma), places its origin in the Early Pliocene, a period marked by substantial environmental and geomorphological changes in Amazonia. This timing aligns closely with divergence estimates reported by Menezes et al. (2010), supporting the hypothesis that *Aotus* originated in the Central Amazon and subsequently diversified in association with the establishment of the modern Amazon River system and ongoing Andean uplift. The estimated age corresponds with the onset of the Pliocene Epoch (5.33–2.58 Ma; Hoorn et al., 2023), one of the most transformative intervals in South America’s geological and ecological history. During this time, continued uplift of the Andes dramatically altered regional climates, reshaped continental drainage patterns, and contributed to the formation of extensive transcontinental river networks. Climatic fluctuations further influenced vegetation dynamics, promoting the expansion of grasslands in some areas and contraction of forested habitats in others, setting the stage for diversification and lineage differentiation in Amazonian taxa such as *Aotus*.

This period of rapid *Aotus* expansion and diversification coincides with major biostratigraphic and paleoenvironmental events, including the dispersal and expansion of herbaceous plant taxa, Late Miocene cooling, Andean uplift, and associated climatic shifts (Hoorn et al., 2010). The transition from the Miocene to the Pliocene (∼5 Ma) marked a shift toward cooler and drier conditions, driving substantial ecosystem changes, including the expansion of grasslands and alterations in forest composition. These environmental changes correspond with rapid diversification in multiple mammalian lineages, such as Caviomorpha rodents (Álvarez et al., 2017), and Didelphidae (Jansa et al., 2013; Castro et al., 2021). Similarly, several Neotropical primate genera experienced accelerated diversification during the Late Miocene, including *Callicebus*, *Cebus, Saguinus*, and *Callithrix* (Springer et al., 2012).

Notably, intense Andean uplift preceded the rapid diversification of *Aotus* by several million years, and vicariance-driven speciation in other primate lineages is documented as early as 11–9 Ma (Alfaro, 2017). This pattern suggests that initial colonization of the Amazon Basin by ancestral *Aotus* may have occurred prior to major Andean orogeny, with subsequent diversification facilitated by climatic shifts, ecological adaptation, and the establishment of new riverine barriers. Collectively, these findings support a model in which geological and climatic changes during the Late Miocene and Early Pliocene were critical drivers of *Aotus* evolution and diversification.

The correspondence between molecular clades and geography supports the recognition of three major evolutionary lineages (i.e., northern, western, and southern) consistent with previous biogeographic models, as described by Plautz et al. (2009). Divergence estimates among the three primary clades overlap within their respective HPD intervals, suggesting a rapid radiation rather than a strictly sequential divergence process. *A. trivirgatus* diverged from the southern clade around 4.1 Ma (95% HPD: 3.7–4.5 Ma), a result consistent with its basal placement in BI analyses and its intermediate position under ML. Such topological variation may reflect incomplete lineage sorting or the effects of rapid speciation during this period. With that said, the timing of diversification into the three clades coincides with major environmental reorganization in Amazonia, potentially reflecting the influence of riverine barriers and habitat shifts on lineage splitting. Geological evidence indicates that the Amazon Basin underwent dramatic drainage reconfiguration during the Neogene, including the transition of the Amazon’s main channel toward eastward discharge, with estimates placing the onset of this transition between ∼10–7 Ma and into the Pliocene. The establishment of a broadly modern eastward-flowing Amazon and its tributary network would have provided both dispersal corridors for arboreal taxa and long-term barrier effects across northern versus southern tributaries. Although the exact timing remains debated, the Late Pliocene to early Pleistocene (∼3-2.5 Ma) appears to mark further refinement of tributary connections, likely influencing the diversification of many Amazonian lineages, including monkeys such as *Aotus*.

The estimated divergence between *A. vociferans* and *A. nancymaae* at 3.4 Ma (95% HPD: 3.0– 3.8 Ma) is congruent with estimates from prior multilocus and mitochondrial analyses (Boubli et al., 2012; Martins-Junior et al., 2022) and likely corresponds to the final stages of Amazon Basin drainage stabilization during the Late Pliocene. Similarly, the split between *A. nigriceps* and the southern *A. azarae* - *A. boliviensis - A. infulatus* complex at approximately 1.5 Ma (95% HPD: 1.2–1.8 Ma) may have been influenced by Pleistocene climatic oscillations, which drove forest contraction and fragmentation throughout southern Amazonia (Ribas et al., 2012).

Within the southern group, the relatively shallow divergence among *A. azarae*, *A. boliviensis*, and *A. infulatus* (1.2–1.6% pairwise divergence) mirrors patterns described by Martins-Junior et al. (2022), who suggested these taxa may represent recently diverged but distinct species. As well, the shallow differentiation observed between *A. miconax* and *A. nancymaae* (0.9% pairwise divergence) and the overlapping ranges of their haplotypes indicate potential historical introgression or ongoing gene flow or incomplete lineage sorting, underscoring the need for expanded sampling and incorporation of nuclear genomic data to resolve their taxonomic status. suggesting incomplete lineage sorting or across their contact zone.

Taken together, the concordance between ML and BI analyses, the overlapping HPD intervals for major diversification events, and the temporal association with key paleoenvironmental transitions all support a model of rapid, river-mediated radiation of *Aotus* during the Pliocene and Pleistocene. These results further strengthen the view that Amazonian River systems, acting in concert with climatic and topographic changes, have been primary drivers of primate diversification in South America (e.g., Ribas et al., 2012; Boubli et al. 2015). Continued integration of genomic, morphological, and ecological datasets will be critical for refining species boundaries within *Aotus* and for testing hypotheses regarding the historical processes underlying its remarkable biogeographic and phenotypic diversity.

Analyses of COI sequences generated from an expanded set of fecal samples, combined with mitogenome-derived and publicly available sequences, corroborate mitogenomic results (Figure 5). The expanded COI dataset also revealed additional fine-scale haplotype structure within several species, providing greater resolution of intraspecific diversity. Multiple haplotypes were identified in *A. trivirgatus*, *A. vociferans*, *A. nancymaae*, *A. nigriceps*, and *A. azarae*, corresponding to populations separated by major Amazonian river barriers. Importantly, the COI analysis allowed differentiation and clarification of the relationship between *A. miconax* and *A. nancymaae* and also facilitated the correction of several previously misidentified sequences, primarily from *A. boliviensis* and *A. lemurinus*. This pattern underscores the role of rivers as persistent barriers shaping genetic structure and suggests that even within currently recognized species, substantial mitochondrial differentiation exists. The distribution of COI haplotypes broadly coincides with recognized Areas of Endemism within the Amazon Basin (Ribas et al., 2022; Helenbrook & Valdez, 2025), reinforcing the hypothesis that *Aotus* diversification has been tightly linked to the spatial configuration of Amazonian landscapes. This concordance highlights the value of incorporating expanded geographic sampling to more fully capture hidden population structure and evolutionary processes operating within the genus, though additional sampling remains necessary, particularly for underrepresented species and understudied regions of the Amazon Basin.

### 4.2 Taxonomy and Conservation Implications

This phylogenomic analysis provide novel resolution of lineage boundaries within *Aotus* and underscore the taxonomic and conservation significance of these findings (Figure 6). Furthermore, analysis of the single-gene COI marker reveals previously unrecognized genetic differentiation, supporting the existence of distinct evolutionary lineages. Beginning in the northern Amazon, a sample previously assigned to *A. vociferans* north of the Japurá River clusters unambiguously with *A. trivirgatus*, expanding the known distribution of this species westward by over 200,000 km² while correspondingly reducing the estimated range of *A. vociferans*. This result highlights the value of comprehensive sampling and molecular verification for refining species ranges and has immediate implications for range-based conservation planning. Furthermore, the presence of two distinct COI haplogroups within *A. trivirigatus*, separated by the Negro River, indicates underlying genetic structure across its range. Similarly, analysis of COI sequences revealed two distinct haplogroups within *Aotus vociferans*, with divergence corresponding geographically to the Rio Napo. Correct delineation of *A. trivirgatus* and *A. vociferans* distribution, taking into account this haplogroup diversity and modified distribution, will inform assessments of population viability and connectivity, guide habitat protection priorities, and help identify potential threats across the northern Amazon. However, additional sampling is necessary from animals of known provenance.

**Figure 6:**
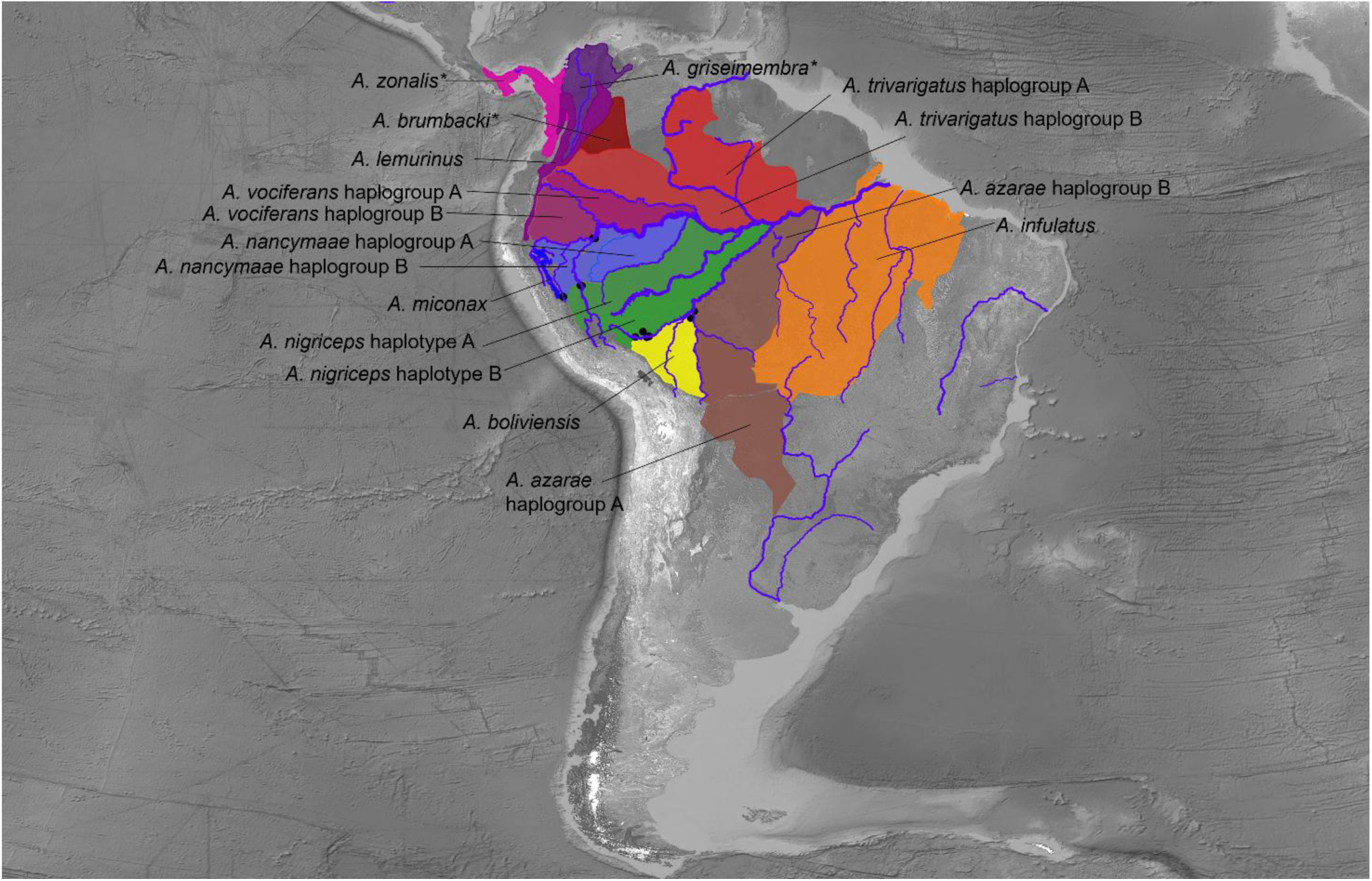
Updated geographic distribution of *Aotus* taxa reflecting results from species delimitation analyses, phylogenomic inference using Bayesian Inference (BI) and Maximum Likelihood (ML), and mitochondrial COI-based phylogenetic analyses. Major river barriers are depicted and correspond to the distribution of all taxa, including newly identified haplogroups. Patterns of lineage divergence across riverine boundaries are consistent across mitogenomic and COI datasets, highlighting the role of Amazonian river systems in shaping *Aotus* diversification.

The conservation status of several *Aotus nigriceps* is likely to be significantly affected by the revised taxonomic and phylogenomic findings presented here. For instance, the range of *A. nigriceps*, historically estimated at approximately ∼1,479,613 km² (Helenbrook and Valdez, 2021) and including areas south of the Madeira River in Brazil, is now revised to roughly 838,295 km² (∼56.7% of the former range) following phylogenomic partitioning. In addition, phylogenomic data revealed two monophyletic haplogroups separated by the Purus River, suggesting either cryptic taxonomic differentiation or deep intraspecific structure. Although pairwise divergence is relatively low (∼0.3%), the existence of geographically structured haplotypes supports recognition of these lineages as provisional management units, warranting monitoring and targeted sampling to determine whether they represent distinct evolutionary significant units (ESUs).

To the east, *A. miconax* and *A. nancymaae* exhibit fairly minimal mitogenomic pairwise divergence (0.99%), and their classification varies depending on the species concept applied. Phylogenetic species delimitation analyses yield mixed results, and the inclusion of a captive-origin *A. miconax* sample underscores the need for additional field-based sampling across the Ucayali–Tingo María region. However, the expanded COI analysis was able to clearly distinguish *A. miconax* from *A. nancymaae*, while simultaneously revealing two distinct haplotypes within *A. nancymaae*. This highlights previously unrecognized genetic structure within *A. nancymaae* and underscores the value of targeted molecular markers, and expanded sampling, for resolving closely related *Aotus* lineages.

Divergence among *A. boliviensis*, *A. infulatus* and *A. azarae* was modest (1.2–1.6%); however, full species-level distinction is supported under the phylogenetic species concept, species delimitation analyses, and single-gene COI data. Samples previously attributed to *A. boliviensis* are now recognized as *A. azarae* haplogroup B (Figure 5), whereas the mitogenomes generated in this study originate from a previously unsampled geographic region and are distinct from both *A. azarae* and *A. infulatus*. Samples obtained from populations in Rondônia, Brazil, exhibit pronounced differentiation in the COI marker, suggesting that formal taxonomic description may be warranted pending additional sampling and morphological characterization. Collectively, these findings justify a cautious, provisional recognition of species status, pending integrative genomic and morphological analyses.

The newly hypothesized taxon currently designated as *Aotus azarae* haplogroup B, occurring south of the Madeira River, inhabits regions experiencing extensive deforestation and habitat degradation in the Brazilian states of Amazonas and Rondônia, with Rondônia exhibiting particularly severe forest loss in recent decades. Given ongoing anthropogenic pressures, including deforestation, wildfires, and climate change, this area in particular is likely to experience elevated rates of population decline. Moreover, the recognition of additional lineages, such as the two monophyletic haplogroups within *A. nigriceps* separated by the Purus River, and the cryptic diversity noted for *A. nancymaae* and *A. vociferans, and A. trivirgatus*, highlights previously unrecognized conservation units that may require targeted protection.

Collectively, these findings underscore the importance of integrating phylogenomic data into conservation planning for *Aotus*. The recognition of three deeply diverged clades, geographically structured haplotypes, and cryptic lineages provides a robust framework for delineating evolutionary significant units (ESUs) and management units across the Amazon Basin. Accurate species and subspecies assignments will improve range maps, inform threat assessments, and guide targeted conservation actions, particularly in regions experiencing unprecedented deforestation and habitat degradation. Moreover, these results highlight the value of combining mitochondrial phylogenomics with morphological, biogeographical, and ecological data to resolve taxonomic uncertainty and safeguard the evolutionary diversity of nocturnal primates in Amazonia.

### 4.3 Fecal metagenomics

In addition to its phylogenetic contributions, this study highlights the value of fecal metagenomics as a noninvasive and cost-efficient method for collecting high-quality molecular data in cryptic primate taxa such as Aotus. For example, shotgun fecal metagenomic sequencing has been used successfully in the banded leaf monkey (*Presbytis femoralis*) to reconstruct full mitochondrial genomes, examine genetic diversity, and simultaneously assess diet and gut parasites from droppings alone (Srivathsan et al. 2016). By enabling broad geographic sampling, without the need for capture or tissue collection, fecal metagenomics allows us to fill previously intractable gaps in *Aotus*’ range, especially in inaccessible Amazonian regions. These data enhance our resolution of lineage relationships, haplotype distributions, and past demographic events in a manner that is both ethically aligned with conservation imperatives and operationally scalable.

Beyond systematics, the broader ecological and health-related applications of fecal metagenomics are numerous. Because all DNA in a fecal sample is sequenced, researchers can infer diet (via plant, insect or other prey DNA), detect parasite communities, and characterize microbiome diversity—each of which can inform evolutionary, ecological and conservation questions. For instance, comparative metagenomic analyses in wild primates show that gut microbial community composition strongly correlates with diet and geography (Srivathsan et al. 2016). In *Aotus*, integrating fecal metagenomic data could shed light on how dietary niche shifts, parasite burdens, or microbiome variation differ among the northern, western and southern clades, thereby linking phylogenetic divergence to ecological adaptation and conservation unit distinctiveness. While the expanded mitochondrial dataset presented here helps fill key geographic and taxonomic gaps, resolving species boundaries, especially among recently diverged taxa, still requires complementary nuclear and ecological data. Nevertheless, this work ushers in a key component of *Aotus* evolutionary history into clearer focus and exemplifies how fecal metagenomics can serve as a foundational platform for integrative primate systematics and conservation.

Continued integrative research remains essential to fully resolve *Aotus* systematics, particularly in undersampled regions north of the Amazon River and extending into Central America. Morphometric analyses, ecological niche characterization, biogeographic surveys, behavioral studies, and expanded nuclear genomic data will be critical for clarifying species boundaries, detecting potential cryptic diversity, and assessing population connectivity. Notably, Areas of Endemism in the Amazon Basin correspond closely with the taxonomic and phylogenomic patterns recovered in this study (Ribas et al., 2022; Helenbrook and Valdez, 2023; Figure 1), providing independent support for the three major clades and several geographically structured lineages. The primary exception is the region northwest of Manaus along the Negro River, for which molecular and morphological data remain lacking, highlighting a key gap for future sampling and conservation assessment.

In total, this study advances our understanding of *Aotus* taxonomy and evolutionary history through comprehensive mitogenomic analyses. By integrating genomic data with biogeographical and ecological insights, the research provides a robust framework for future studies on Neotropical primate conservation and management. Continued research efforts, particularly in underrepresented regions like Central America and the northern Amazon, are essential for further unraveling the complexities of *Aotus* evolution and ensuring effective conservation strategies. Increased taxonomic resolution and the designation of new conservation management groups could conceivably mean that some of these taxa face far more dire scenarios than current analysis would suggest.

## Acknowledgements

Special thanks to Jeissy Evelyn Huamani and Jessica A. Suarez for their contribution in the field.

